## Supplemental Data for "Multiple Guidance Mechanisms Control Axon Growth to Generate Precise T-shaped Bifurcation during Dorsal Funiculus Development in the Spinal Cord"

#### Supplemental Figure Legends

**Fig. S1 *Ntn1* expression in mouse embryonic spinal cord and additional wholemount staining in E10.5 embryos**.

(**A-B**) Dark field images of *in situ* hybridization using $^{35}$S-labeled mRNA probes for *Ntn1* in cross sections of E10.5 and E11.5 mouse embryos. *Ntn1* expression is mainly confined to the inside of the spinal cord, including the floor plate (FP), the ventricular zone, and the lateral domain in the dorsal horn adjacent to the DREZ (arrow). Asterisks (*) label the DRG.

(**C-J**) Whole mount NF staining using HRP-based colorimetric substrates in E10.5 WT (C-D,G-H) or *Ntn1*$^{\beta/\beta}$ (E-F, I-J) embryos (n=5). BA/BB-cleared embryos are viewed from the lateral side of the forelimb (C-F) or hindlimb (G-J) region. Asterisks (*) label the DRG, arrows point to the front (yellow) or the back (white) dorsal funiculus, and misprojections (black). Images in D,F,H,J show enlarged views of the boxed region of a single DRG in C,E,G,I.

Bars: 100 μm.

**Fig. S2 Traditional wholemount NF immunostaining reveals the loss of the dorsal funiculus in *Slit1*$^{-/-}$*;Slit2*$^{-/-}$*;Ntn1*$^{\beta/\beta}$ triple mutants**

(**A-B**) HRP-based neurofilament staining is viewed from the dorsolateral side of E10.5 embryos in the forelimb (A-B) or the hindlimb (C-D) region. Note the difference in labeled axons at the front (yellow arrowheads) or the back (white arrowheads) dorsal funiculi after they grow out from the DRG (*) between *Slit1*$^{-/-}$*;Slit2*$^{-/-}$*;Ntn1*$^{\beta/\beta}$ mutants (B,D) and littermate controls (A,C) (*n*=2).

**Fig. S3 *DCC* expression in mouse embryonic spinal cord**.

Dark field images of in situ hybridization using $^{35}$S-labeled mRNA probes in cross sections (16 μm) of mouse spinal cords from E10.5 and E11.5 embryos. *DCC* is expressed mainly inside the spinal cord and weakly in the DRG (*). FP: floor plate.

**Fig. S4 Cartoon illustration of the different guidance roles of Slit and Ntn1 during bifurcation**.

### Fig. S1

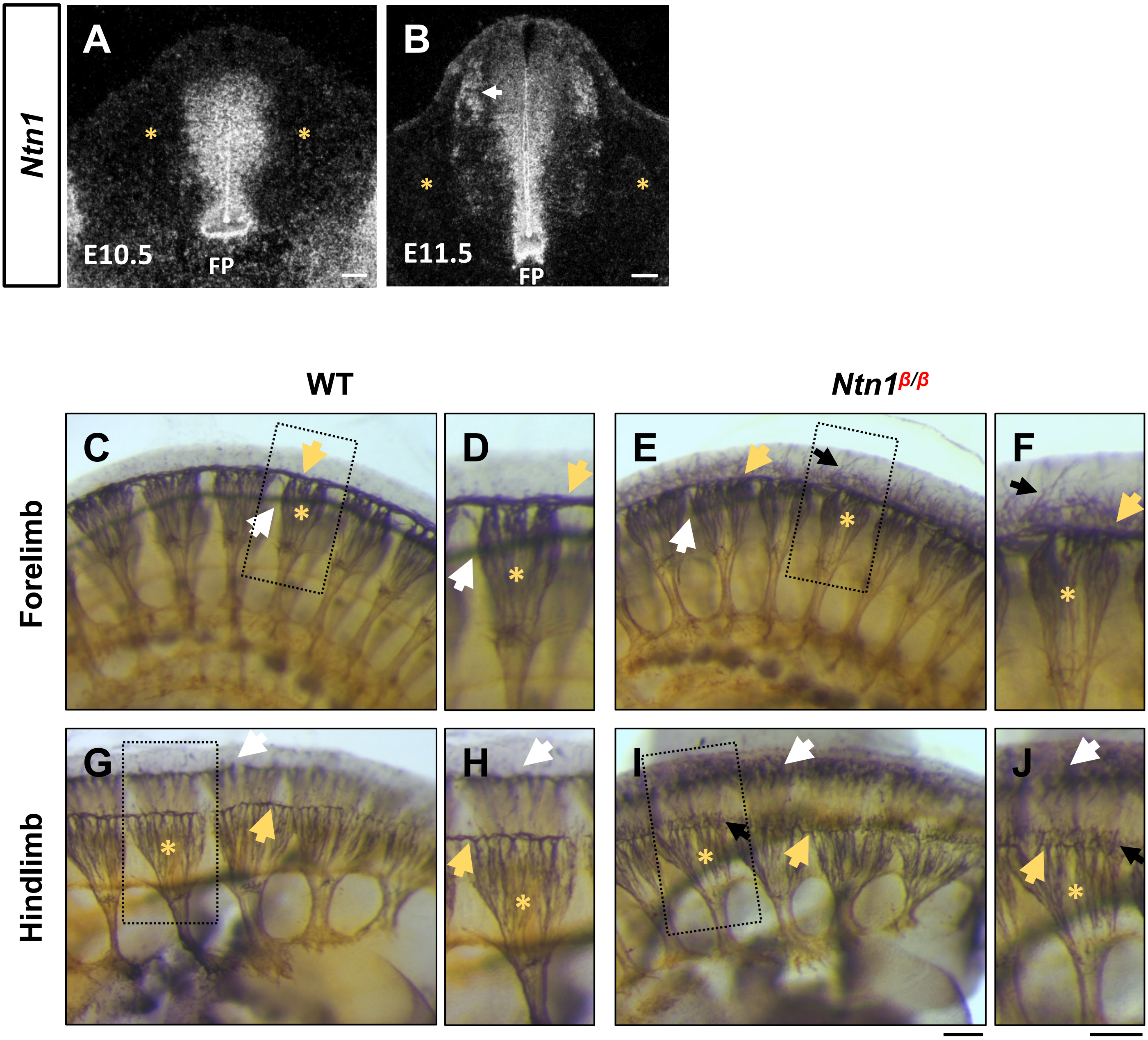

Ntn1
A
B
E10.5
FP
E11.5
FP
WT
$Ntn1^{\beta/\beta}$
Forelimb
Hindlimb
C
D
E
F
G
H
I
J

Fig. S2

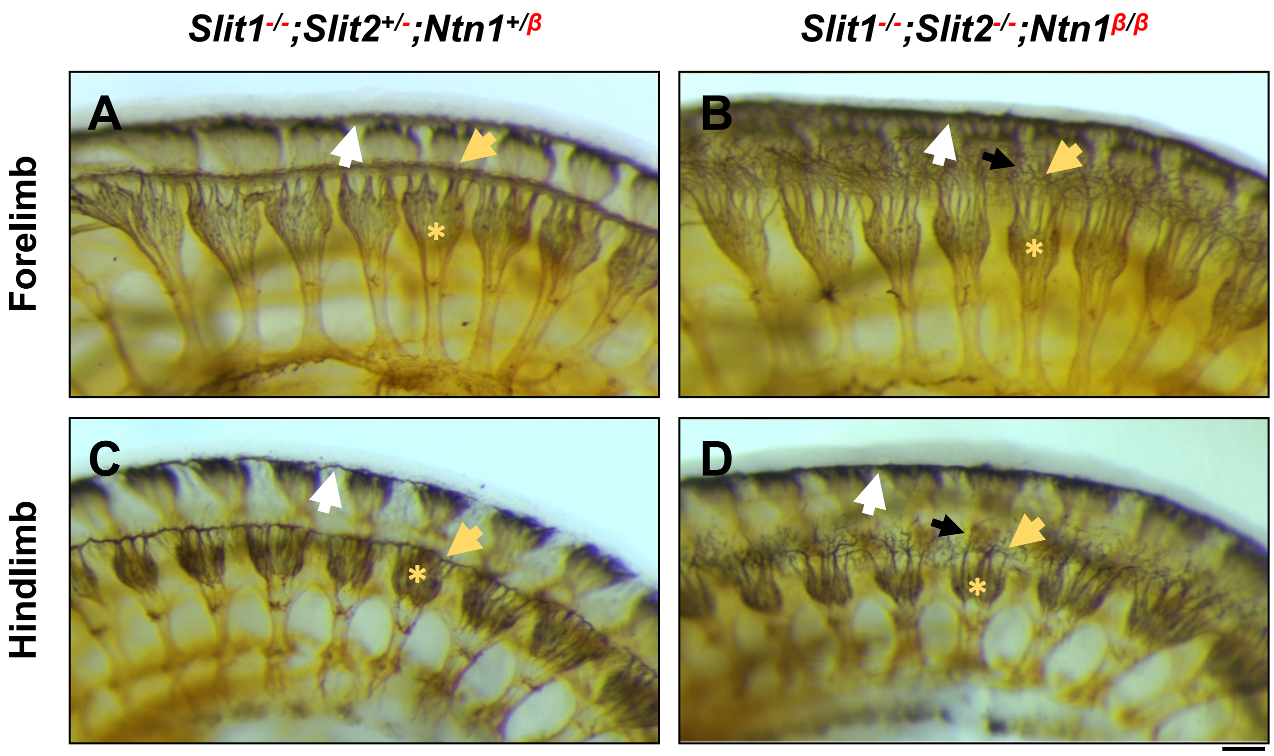

Slit1-/-;Slit2+/-;Ntn1+/β
Slit1-/-;Slit2-/-;Ntn1β/β
Forelimb
Hindlimb
A
B
C
D

Fig. S3

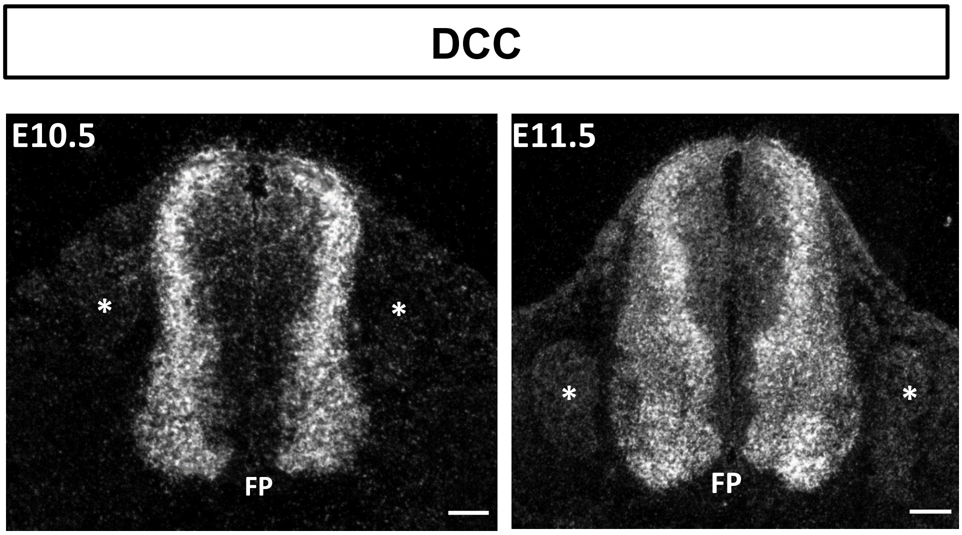

Fig. S4

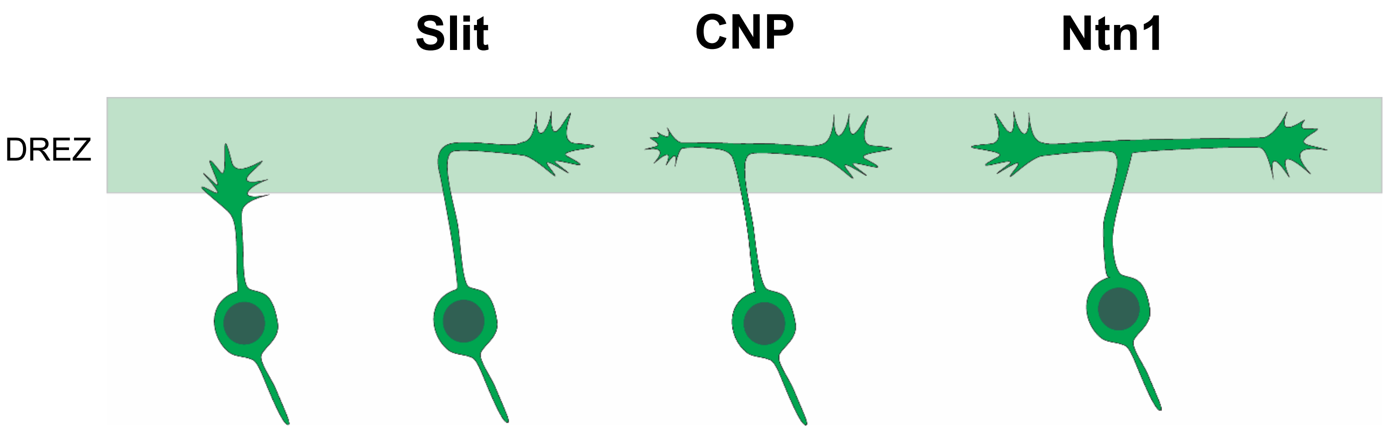
